# High-throughput deep tissue two-photon microscopy at kilohertz frame rates

**DOI:** 10.1101/2022.10.24.513414

**Authors:** Sheng Xiao, John T. Giblin, David A. Boas, Jerome Mertz

## Abstract

Strategies to image biological phenomena at millisecond time scales are generally technically challenging and suffer from compromises between imaging field-of-view, depth penetration and excitation efficiency in thick tissue. We present a simple and cost-effective solution that enables a conventional video-rate two-photon microscope (2PM) to perform 2D scanning at kilohertz frame rates, while preserving all the benefits of standard 2PM, which we demonstrate by imaging neurovascular dynamics in mouse brains.

## Main

Metabolic or cellular activity, such as electrical signaling^1^, cerebral blood flow^2^, glutamate release^3^, etc., can occur on time scales of milliseconds. The development of optical microscopes capable of monitoring such fast activity has been a topic of intense research^4^. Though widefield microscopes can attain the required speeds when equipped with modern sCMOS cameras, they fail to provide inherent background rejection, limiting their ability to image thick or densely labeled tissue^5^. On the other hand, conventional laser scanning microscopes (LSMs) such as confocal or multiphoton microscopes, though they provide inherent background rejection and thus improved contrast, are limited by the speed of their scanning mechanisms.

In most cases, the scanning mechanism of a LSM involves the mechanical movement of mirrors, and is thus limited by inertia. For example, galvanometer-mounted mirrors, even when resonant, typically offer only video rate imaging (~30 Hz). Faster imaging can be achieved by reducing the size of the scanning mirrors (e.g. high-facet-count polygonal scanning^6^), but comes at the cost of reduced throughput, as manifested by a reduced field-of-view (FOV) or resolution^7^. Alternatively, faster imaging can be achieved by parallelization using multi-focus scanning and array detection with a camera or multi-pixel detector^8, 9^. However, tissue scattering in the detection path leads to crosstalk ambiguity between nearby foci which limits depth penetration. Such ambiguity can be alleviated with the use of a priori sample information, for example combined with compressed sensing^10^, but this too is restricted to sparse or pre-mapped samples.

Still another strategy for fast imaging makes use of scanning mechanisms that are inertia free, for example using acousto-optic deflectors^11–13^ or passive pulse splitting^14^. Since both approaches exploit single-point scanning and single-pixel detection, they are advantageous for maintaining high contrast at large tissue depths^15^. However, at kilohertz frame rates these scanning mechanisms exhibit limited throughput, where only a few cells can be imaged simultaneously. Moreover, the requirement for specialized instrumentation (e.g. a low repetition-rate high pulse energy laser^14^) impose practical constraints that have hindered their general adoption. To date, the availability of two-photon systems that provide the combination of high-contrast, high-speed, and large FOV has remained elusive.

Here we present a kilohertz two-photon microscope (2PM) for high-throughput deep tissue imaging. Our technique consists in modifying an entirely conventional 2PM (80 MHz laser, video-rate resonant-linear galvanometer scanning, PMT detection, etc.) with the addition of a simple device called a scan multiplier unit (SMU)^16^ comprising a scan lens, a microlens array, and a mirror. The SMU exploits a double-pass geometry that has the effect of multiplying the scanner rate without having to modify the operation of the scanner itself, while at the same time doubling scanner throughput. Because the SMU contains no moving parts, the multiplication is inertia-free, with a multiplication factor determined by the number of lenslets in the microlens array. Moreover, since the laser beam is not split into multiple beamlets, SMU-2PM maintains high excitation efficiency allowing the use of a common 80 MHz repetition-rate laser with conventional single-detector signal acquisition at standard sampling rates (here 80 MHz).

The basic principle of a SMU was introduced in Ref.^16^. Here, to achieve kilohertz-rate scanning, we used a 16 × 1 lenslet array in our SMU, which multiplied an 8 kHz resonant scanner scan rate to 256 kHz for fast-axis scanning [SMU1 in Fig. 1(a)]. The addition of a 1 kHz slow-axis galvanometer then enabled 2D raster scanning at a frame rate of 1 kHz over a FOV of 200 × 200 μm with a 16× objective. The spatial resolution of our microscope is simultaneously affected by the optical point-spread-function (PSF) (Supplementary Fig. 3, 4), laser pulse sampling density, and the accuracy of our image registration (see Methods, Supplementary Fig. 5, 6, 7). Overall, this was measured to be 2.2± 0.8, 1.9 ± 0.7, 8.2± 3.0 μm [mean ± S.D.; full-with-at-half-maximum values (FWHM) obtained by Gaussian fitting] along the x, y, and z axes respectively. We note that different scan rates can be readily achieved with different lenslet counts. To enhance the usability of our microscope, we included a second SMU whose lenslet array was replaced by a single lens [SMU2 in Fig. 1(a)], enabling the microscope to revert to its original video-rate scan configuration, though with a doubled fast-axis scan angle. Switching between SMU1 and SMU2 with a flip mirror thus allowed a convenient toggling between different FOV sizes and frame rates, essentially providing a zoom mechanism to situate the kilohertz FOV.

**Figure 1.**
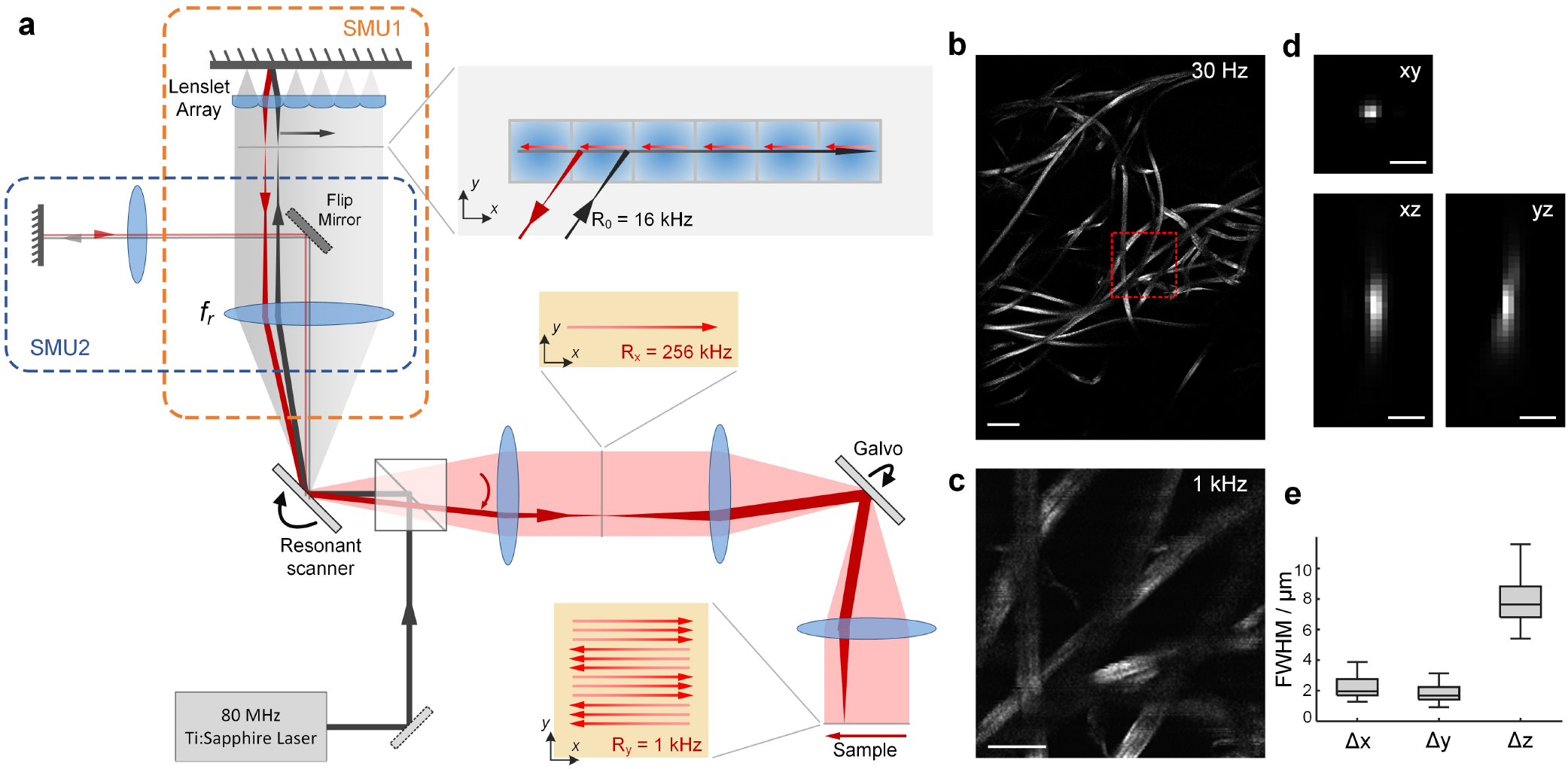
SMU-based 2PM for dual video-rate/kilohertz frame rates imaging. (a) Schematic of microscope. (b) Conventional video-rate two-photon imaging over a large FOV using SMU2. Scale bar 100 μm. (c) Kilohertz two-photon imaging over a 200 × 200 μm FOV using SMU1. Scale bar 40 μm. (d) Example PSF measured with SMU1 using 100 nm fluorescent beads. Scale bar 5 μm. (e) Lateral and axial FWHM of PSFs (n = 114) measured across entire FOV using 100 nm fluorescent beads. FWHM obtained by fitting the intensity profiles to a Gaussian. Box plots: box, 25th to 75th percentiles; whiskers, 1.5× interquartile range from the 25th and 75th percentiles; middle horizontal line, median.

The ability to monitor fast, complex dynamics over large FOVs is critical for many imaging applications. An example application that highlights the advantages of SMU-based 2PM is the study of neurovascular coupling. We begin here by imaging cerebral blood flow in awake mice using inverse fluorescence contrast of red blood cells (RBCs). Blood flow rates are determined from the dominant orientation of RBC kymographs, which, traditionally, can be measured on single blood vessels by rapid 1D line scanning^2^. More recently, FACED-based 2PM^17^ was shown to enable kilohertz imaging of a few vessel segments at a time. Here, by virtue of the high throughput of SMU scanning, we routinely imaged more than 10 vessel segments at a time, allowing connections between multiple capillaries, venules, and arteries to be monitored simultaneously, and a wide diversity of RBC flow speeds to be quantified [Fig. 2(a-d), Supplementary Fig. 8, 9, Supplementary Video 1-3]. Within the same FOV, we observed both positive (vessels 3-5) and negative (vessels 5, 6) velocity correlations across connecting vessels, and positive correlations across distant vessels (vessels 1-2 to 6-11) [Fig. 2(d)]. In larger vessels, velocity profiles indicative of laminar flow could be spatially resolved (Supplementary Fig. 9, Supplementary Video 2). We note that at reduced acquisition speeds of 30 Hz, capillary flow could not be quantified (Supplementary Fig. 11).

**Figure 2.**
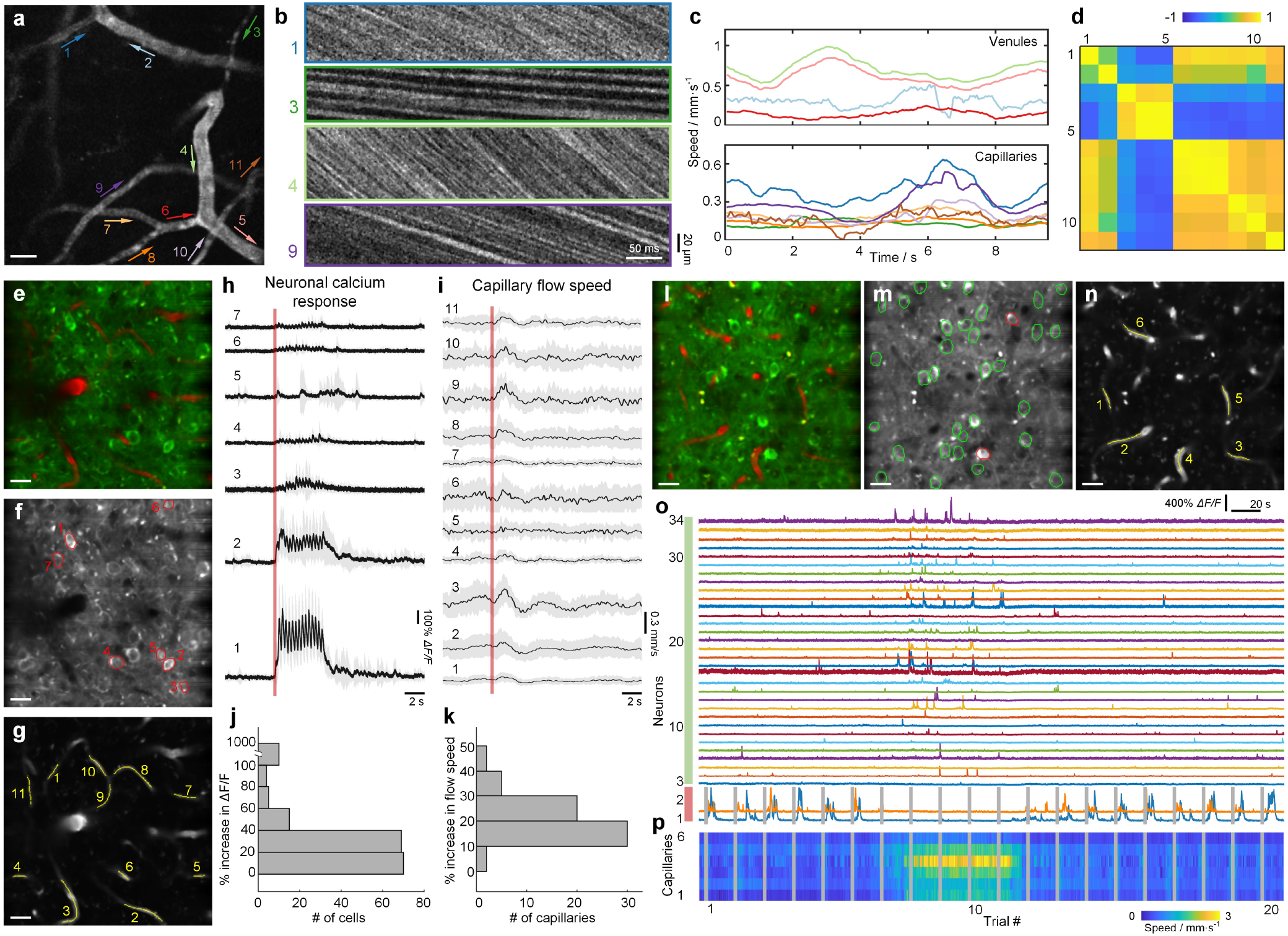
SMU-2PM enables high-throughput imaging of fast, complex biological dynamics in vivo. (a) Brain vascular network imaged at 1 kHz frame rate over a 200 × 200 μm FOV. (b) Kymographs of 4 selected vessel segments labeled in (a). (c) Flow speeds of 11 vessel segments measured over a 10 s recording. (d) Cross-correlation coefficients of the velocity profiles for all vessel segments shown in (a). (e) Composite of simultaneous GCaMP (green) and Texas Red (red) imaging in vivo. (f) Averaged image of GCaMP6 fluorescence. 7 neurons (outlined in red) within the FOV showed response evoked by whisker stimulation. (g) Averaged image of Texas Red fluorescence. 11 capillaries (outlined in yellow) within the FOV showed increased flow speed during whisker stimulation. (h,i) Trial-averaged (n = 20 trials) calcium signals and capillary flow speed for 7 neurons and 11 capillaries that showed response to whisker air-puff stimuli, as outlined in (f,g). Grey shaded areas represent ±1 standard deviation. Red vertical lines indicate the onset of whisker stimulation. Each trial consisted of 5 s baseline recording, followed by 5 s whisker air-puff stimulation (3 Hz rate), followed by 10 s post-stimulation recording. (j) Maximum percentage increase in trial-averaged Δ*F/F* during the 5 s stimulation period compared to the baseline Δ*F/F*. Histogram of 173 neurons from 5 different FOVs. (k) Maximum percentage increase in trial-averaged capillary flow speed during the 5 s stimulation period compared to the baseline flow speed. Histogram of 59 capillaries from the same 5 FOVs used in (j). (l-n) Simultaneous imaging of GCaMP6 (m) and Texas Red (n) from another FOV. 2 neurons showing whisker-evoked response are outlined in red, 32 neurons not showing responses are outlined in green. 6 capillaries with quantified flow speeds are outlined in yellow. (o) Calcium traces for all 34 neurons. Each trace is assembled from 20 individual trials (20 s per trial). (p) Corresponding flow speed from 6 capillaries shown in (n), temporally aligned with (o). Vertical gray lines in (o,p) show the onset of each whisker air-puff stimulation. Imaging depth for (a-d) was approximately 50 μm below brain surface, with 50 mW excitation power. Imaging depths for (e-p) were 120-180 μm, with 100 mW excitation power. Scale bars in (a), (e-g), (l-n) are 20 μm.

The study of neurovascular coupling requires dual imaging of both blood flow and neuronal activity at network level. On a 2PM, this is possible at low speeds (< 100 Hz) by utilizing arbitrary scan patterns^18^. Improving temporal resolution requires the combination of a 2PM with an alternative high-speed imaging modality^19^, which comes at the expense of spatial resolution. Here, simply by adding a second detection channel capable of monitoring GCaMP6f fluorescence, capillary blood flow can be correlated with local calcium activities even in dense populations of neurons (>50 per FOV, Supplementary Fig. 10, Supplementary Video 4). Under whisker air-puff stimulation (n = 20 trials per FOV), we observed a reliable calcium response from a subset of neurons [Fig. 2(h)], and a simultaneous increase in flow speeds across the capillary network [Fig. 2(i)]. Across 5 different FOVs, 10 out of 173 neurons showed strong responses to whisker stimuli (> 100% maximum increase in trial-averaged Δ*F/F*), and 59 capillaries showed an average increase of 20% in maximum flow speeds [Fig. 2(j,k)]. In one FOV, we observed a suppression of whisker-evoked responses in 2 responding neurons across multiple trials [neuron 1 and 2 in Fig. 2(o), trial 8 - 11], coinciding with an increase in activity observed elsewhere [neuron 3-34 in Fig. 2(o)], as well as a large increase in flow speeds (average 158%) of all 6 quantified capillaries within the FOV [Fig. 2(p)]. Such a diversity of vascular network flow and its coupling to different subsets of neurons could not have been revealed without the speed and throughput capacity of SMU-2PM imaging. To our knowledge, this is the first observation of neurovascular coupling throughout an extended FOV with microcapillary resolution and at kilohertz frame rates.

We emphasize that a fundamental advantage of SMU-2PM over alternative multifocus or line scanning approaches is that it benefits from the well-known depth penetration advantages of conventional single-point-scanning 2PM, namely increased signal contrast and compatibility with large-area single-pixel detection^15^. In mouse brains expressing GCaMP6f, we were able to consistently reach penetration depths > 500 μm (n = 4 FOVs in 2 mice) and observe calcium transients from individual neurons, as illustrated in Fig. 3 and Supplementary Video 5-8.

**Figure 3.**
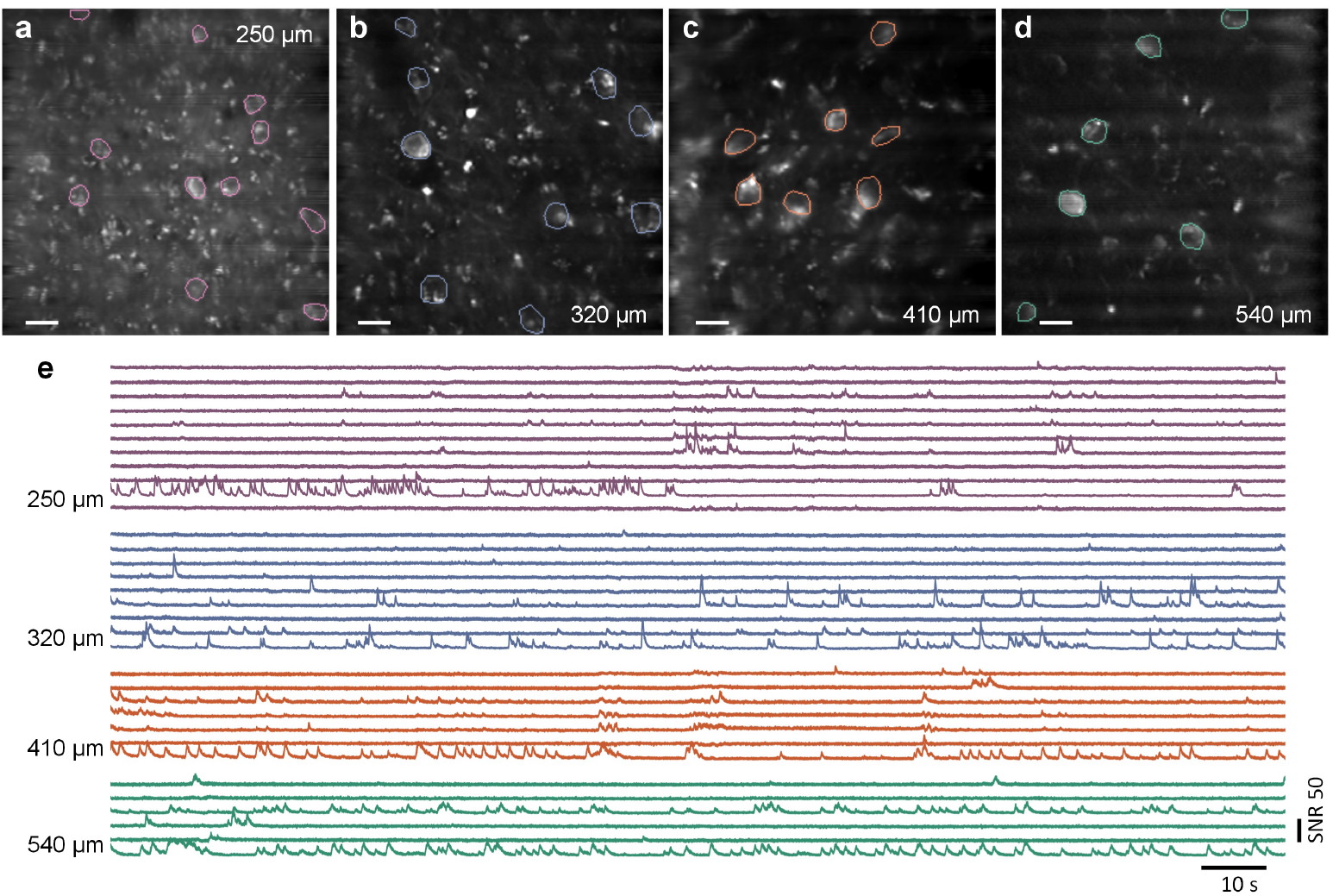
SMU-2PM enables deep tissue imaging of neural activity in vivo. (a-d) Averaged GCaMP images acquired at different depths with 1 kHz frame rates. From (a) - (d), imaging depths were 250 μm, 320 μm, 410 μm, 540 μm, excitation power 160 mW, 220 mW, 220 mW, 250 mW; recording duration 3 minutes. Scale bars are 20 μm. (e) Corresponding calcium traces for outlined neurons in (a-d).

In summary, we have developed a new SMU-based 2PM for kilohertz frame-rate imaging. Our approach involves the addition of passive off-the-shelf optical components to a conventional 2PM (Supplementary Table 1), while also preserving the capacity of the microscope to perform standard video-rate imaging (via SMU2). Different SMU configurations allow easy toggling between different scan rates and FOVs. Importantly, these advantages do not come with reduced excitation efficiency, imaging throughput or penetration depth. As shown here, with 200 × 200 μm FOV and > 500 μm penetration depth, our system enables high throughput imaging of both neural activity and hemodynamics at kilohertz rates in the mouse brain. The cost-effectiveness and practicality of SMU-2PM make it an attractive tool for high-speed imaging in thick tissue.

## Methods

### Optical system design

The schematic for our dual video-rate/kilohertz two-photon microscope is shown in Supplementary Fig. 1. The output of a Ti:Sapphire laser (Coherent Chameleon Ultra II, 3.5 W output power, 920 nm wavelength, 80 MHz repetition rate, 140 fs pulse width) was expanded 3 × using a 4f system (*f*_1_ and *f*_2_). To compensate for temporal dispersion induced by the optical elements, a single-prism pulse compressor^20^ was used to pre-chirp the pulse. The laser beam was scanned by a resonant scanner (Cambridge Technology CRS8kHZ) and then directed into either of the two SMUs selected by a flip mirror. For 1 kHz frame rate imaging, the SMU1 consisted of a scan lens (Thorlabs LSM54-850), a 16 × 1 lenslet array (Amus GmbH APO-GT-P1500-R3.25; p = 1.5 mm, f = 3.98 mm), and a planar mirror. For conventional video-rate imaging (SMU2), the lenslet array was replaced with a single *f*_11_ = 75 mm lens, such that the fast-axis scan rate reverted to 16 kHz. A combination of a polarizing beamsplitter (Thorlabs CCM1-PBS252), a half-wave plate (Thorlabs AHWP05M-980), and a quarter-wave plate (Thorlabs AQWP10M-980) was used to separate the output rescanned beam from the input beam into the SMUs, and also to adjust the post-objective excitation power. After separation, the output of the SMU was collected by a scan lens *f*_3_, which, together with a lens *f*_4_, imaged the resonant scanner surface onto a linear galvanometer (Cambridge Technology 6215H) for slow-axis scanning. The 2D scanned beam was then imaged onto the back aperture of the objective by another 4f system (*f*_5_ and *f*_6_), and focused by the objective (Nikon LWD 16×/0.8NA W) into the sample.

The resulting fluorescence signal was separated from the excitation beam by a dichromatic mirror (DM1, Thorlabs DMLP650L), and directed onto two PMTs for simultaneous dual-color detection. The dichromatic mirror (DM2), and emission filters (Em1, Em2) were chosen according to the fluorescent indicators used. For single channel imaging of Fluorescein-Dextran, Texas Red-Dextran or GCaMP fluorescence, only PMT1 (Hamamatsu Photonics H7422PA-40) with a 680 nm shortpass filter (Em1, Chroma Technology ET680sp-2p8) was used for detection, and the dichromatic mirror Dm2 was absent. For simultaneous calcium and blood-flow imaging, a 567 nm short pass dichromatic mirror (Thorlabs DMSP567R) was used to separate the two channels, with GCaMP fluorescence detected by PMT1 through a 520/70 nm bandpass emission filter (Em1, Edmund Optics 87-735), and Texas Red fluorescence detected by PMT2 (Hamamatsu Photonics R11322U-40-01) through a 680 nm short pass filter (Em2, Chroma Technology ET680sp-2p8).

To help situate the imaging area before each two-photon imaging session, an additional widefield imaging path with 470 nm LED excitation was integrated into the fluorescence detection path, which could be selected with a flip mirror.

### Signal collection and hardware control

The electronics for signal collection and microscope control are shown in Supplementary Fig. 2. Signals from the two PMTs were amplified by a transimpedance amplifier (TPA1, Femto DHPCA-100; TPA2, Femto HCA-100M-50K-C) and lowpass filtered (Mini-Circuits BLP-50+) to 48 MHz to avoid aliasing upon digitization. For each channel, the signal was then split (Mini-Circuits ZFRSC-2050+) into two different digitizers for either 1 kHz or 30 Hz frame rate imaging. Switching between the two imaging modes involved simply switching the flip mirror and running the corresponding Matlab script, without restarting any microscope hardware or software.

For 1 kHz frame rate imaging (SMU1), digitization was performed by an Alazar ATS 9440 digitizer at 80 MHz sampling rate, with all data streamed to disk for postprocessing. A multifunction I/O device (DAQ, National Instrument PXIe-6341) was used to control the resonant scanner scan angle, linear galvanometer scan position, and the acquisition start of the digitizer. Specifically, the linear galvanometer position was controlled by a 1 kHz smoothed triangular wave defined by y = arcsin[(1 – ξ) sin(2*πt/f*)]/*π*, where *ξ*= 0.05 is the smoothing factor, *f* = 1 kHz is the frame rate, and *t* is time. Both the digitization and DAQ clock were synchronized to the 8 kHz resonant scanner frequency using a clock generator (Analog Devices AD9516). A custom Matlab script was used to control the synchronization between the scanner and data acquisition.

For video-rate imaging (SMU2), all microscope hardware, data readout and image reconstruction were controlled by ScanImage software^21^. Digitization was performed by a National Instruments 5771 digitizer, and processed by a National Instrument 7972 FPGA. Scanners were controlled by the same National Instrument DAQ PXIe-6341.

Note that for added flexibility we used two separate sets of detection electronics, benefiting from the fact that the Alazar digitizer allows direct streaming of raw data to the disk at 1 kHz frames rates, which allowed us to evaluate different post-processing algorithms for image reconstruction. In principle the detection electronics can be simplified by using a single digitizer/FPGA set, since both imaging modes make use of the same 80 MHz sampling rate.

### System calibration and image reconstruction

A knowledge of the laser spot position at all times is crucial for the accurate reconstruction of an image. For video-rate imaging (SMU2), our system is identical to a conventional 2PM where image reconstruction relies on the assumption of sinusoidal motion of the resonant scanner along the fast-axis (here with scan amplitude doubled by SMU2), and linear motion from the galvanometer along the slow-axis. The resonant scanner phase was adjusted manually such that no residual sawtooth pattern was apparent along the fast-axis.

When operating at kilohertz frame rates (SMU1), we found that a simple geometric model for the laser spot position along the fast-axis was inadequate because of small alignment and telecentricity errors in the SMU. We thus experimentally calibrated the fast-axis rescan positions by physically translating a single fluorescent bead along the fast-axis, while recording the time when the bead signal appeared within a complete resonant scanner line period. The trajectory was then fitted to a smoothed spline function, obtaining a conversion function between resonant scanner physical scan angle and fast-axis rescan position [Supplementary Fig. 5(a)]. The trajectory of the slow-axis galvanometer when operated at 1 kHz also tended to deviate from the command signal due to physical inertia. We similarly calibrated its position over a 1 ms frame time [Supplementary Fig. 5(b)]. With a knowledge of both the slow- and fast-axis scan positions, we were able to create a 2D scatter plot of laser pulse locations within each frame [Supplementary Fig. 5(d)]. A 2D image based on regularly spaced Cartesian coordinates of size 256 × 256 pixels was then reconstructed using the Matlab *scatteredInterpolant* function with ‘natural’ interpolation.

Because of the nonlinear speed of the resonant scanner, the FOV was not uniformly sampled. While this may not be a problem for video-rate imaging where each pixel can be averaged over multiple laser pulses, it became a problem at 1 kHz rate imaging where the pixel rate was so high that it became comparable to the laser repetition rate, leading to situations where a pixel might not receive even a single laser pulse [Supplementary Fig. 6(d,e)]. This issue was alleviated by adjusting the phase of the linear galvanometer such that the entire FOV was approximately covered during a frame [Supplementary Fig. 5(d,e)], leading to an average laser pulse sampling density of 1.03 ± 0.49 (mean ± S.D.) laser pulses per μm^2^. Alternatively, a linear speed polygonal scanner for fast-axis scanning could also be considered here.

Another complication when performing image reconstruction arose from the drifting in amplitude and phase of the resonant scanner that occurred over time. In the fitted model of the rescan position, we accounted for such drifting between the initial calibration and the actual imaging session by treating the resonant scanner amplitude and phase as an adjustable parameter. Similar to the scanner phase adjustment in a traditional video-rate 2PM, these parameters were manually adjusted to minimize any apparent artifacts in the final reconstructed image (Supplementary Fig. 7). After reconstruction of each frame, the videos were motion corrected using NoRMCorre^22^.

Finally, owing to a non-uniform distribution of intensities across different rescanned lines produced by the SMU, our reconstructed images exhibited minor horizontal striping artifacts. We corrected for these by acquiring a reference image *I_r_* of a uniform fluorescent sample, which we used to normalize the recorded intensity by *I* = *I_raw_*/[*I_r_*/max(*I_r_*) + *η*], where *I_raw_* is the actual fluorescence intensity recorded at each pixel and *η* ∈ [0.2,0.5] is a regularization factor to avoid division by zero.

Throughout this manuscript, averaged fluorescence images are contrast adjusted for better visualization. Supplementary videos are the raw captured frames at 1 kHz.

### Animal preparation and imaging

All animal procedures were approved by the Boston University Institutional Animal Care and Use Committee and were conducted following the Guide for the Care and Use of Laboratory Animals. Mouse strain C57BL/6J (Jackson Laboratory) was used for blood flow only imaging, and C57BL/6J-Tg(Thy1-GCaMP6f)GP5.17Dkim/J (Jackson Laboratory) was used for all calcium related imaging. As described previously^23^, animals were first anesthetized with isoflurane (3% induction, 1-1.5% maintenance). Animals were placed in a stereotaxic frame and the depth of anesthesia was assessed from the respiration rate and toe pinches. A heating blanket was used to maintain the body temperature at 37°C. A craniotomy over both hemispheres was performed with care to keep the dura intact, and the brain surface was covered with a curved glass window^24^ (Lab Maker GmbH) sealed with dental acrylic. A custom ring-shaped head bar made of titanium was secured to the skull with dental acrylic and low viscosity cyanoacrylate (Loctite 401 Instant Adhesive). Animals were given at least two weeks for recovery from the craniotomy procedure prior to habituation to head fixation. For habituation, mice were head-fixed daily in a custom cradle for increasingly longer durations, starting with 15 min/day and until they could be head fixed for 90 minutes. Mice were given sweetened condensed milk as a reward during habituation, and immediately removed if they showed extended signs of discomfort or distress.

For the imaging of cortical vasculature, animals were briefly placed under anesthesia and retro-orbitally injected with 0.03-0.05 mL of Texas Red Dextran (150 kDa, 5% in PBS) or Fluorescein-Dextran (150 kDa, 5% in PBS). During imaging, the animals were head-fixed under the microscope and constrained onto a goniometric platform.

For the imaging of neurovascular coupling dynamics, whiskers on the left side of the animal were deflected by 60 ms duration air-puffs at 3 Hz frequency (Parker Hannifin Corp. Picospritzer III). Imaging was performed near the somatosensory cortex in the right hemisphere of the brain at depths ranging from 120 to 180 μm. For each FOV, we performed 20 trials at 30 s intervals. The recording for each trial lasted 20 s in total, consisting of 5 s recording of a baseline signal, 5 s recording with whisker air-puff stimulus, and 10 s recording of a recovery period.

### Blood flow data analysis

Blood-vessel segments were manually traced to generate the corresponding kymographs. Kymographs were then spatially de-meaned to remove intensity variations along the vessels. An additional median filter of window 3 × 3 pixels was applied to kymographs. To calculate flow speed, we used existing techniques based on either iterative Radon transforms^25^ or image cross-correlation^26^, with a time bin varying from 80 to 500 ms. Cross-correlation matrices of velocity profiles were generated using the Matlab *corrcoef* function.

To evaluate flux rate in capillaries, elliptical regions-of-interest (ROIs) were manually chosen along capillary cross-sections. Temporal intensity fluctuation profiles were created by integrating the pixel values within the ROIs across frames, and RBCs were identified by intensity dips. To count individual RBCs, we applied a lowpass Gaussian filter to the intensity profiles to remove excessive noise, and used the Matlab *findpeaks* function to count the number of intensity dips. The final flux rate was averaged over a 1 s window.

To analyze flow speed changes during whisker air-puff stimulation, we averaged velocity profiles across 20 individual trails for each capillary. The baseline speed *v_b_* was determined from the average flow speed over the first 5 s recording before stimulation, and the maximum flow speed *v_max_* was determined within the following 5 s air-puff stimulation period. The percentage increase in flow speed during stimulation was calculated from (*v_max_* – *v_b_*)/*v_b_*.

### Calcium data analysis

ROIs were manually selected for each neuron based on the time-averaged images, and calcium signals *F* were extracted by integrating all pixel values within the ROIs and median filtering over 10 ms time windows. Calcium trace baselines *F*_b_ were estimated as the average signal intensity of all pixels with intensity below 70th percentile. Trace noise *σ_F_* was estimated as the standard deviation of *F* after Gaussian high-pass filtering (50 ms standard deviation). The Δ*F/F* trace was calculated as Δ*F/F* = (*F* – *F_b_*)/*F_b_*, and the SNR trace was calculated as *SNR* = (*F* – *F_b_*)/*σ_F_*. A comparison of traces with and without median filtering is shown in Supplementary Fig. 12. Percentage changes in Δ*F/F* during whisker air-puff stimulation were calculated similarly to the flow speed changes by comparing the maximum Δ*F/F* obtained during the 5 s stimulation period with the average Δ*F/F* obtained during the preceding 5 s baseline period.

## Acknowledgements

The project was funded by the National Institutes of Health (R01NS116139).

## Author contributions statement

S.X. conceived of, designed and built the SMU-2PM. J.T.G. prepared the experimental animals. S.X. and J.T.G performed the imaging experiments. S.X. analyzed the data. S.X. and J.M wrote the manuscript with contributions from J.T.G. and D.A.B. J.M. and D.A.B. supervised the project.

## Additional information

S.X. and J.M. filed an international patent application No. PCT/US2022/037125 associated with the design and use of a SMU.

## Supplementary Information

**Supplementary Figure 1.**
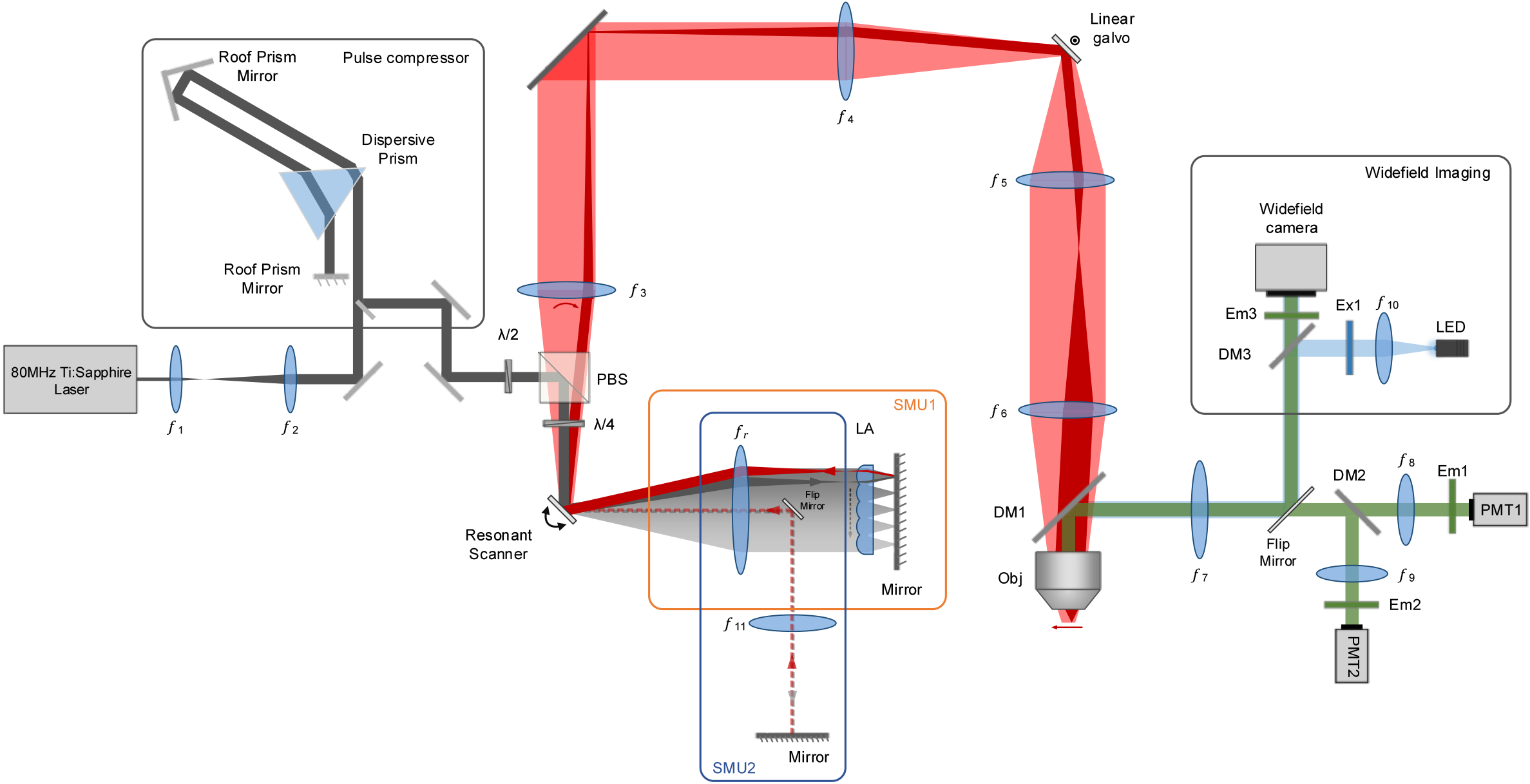
Schematic of the microscope. Components are listed in Table 1.

**Supplementary Figure 2.**
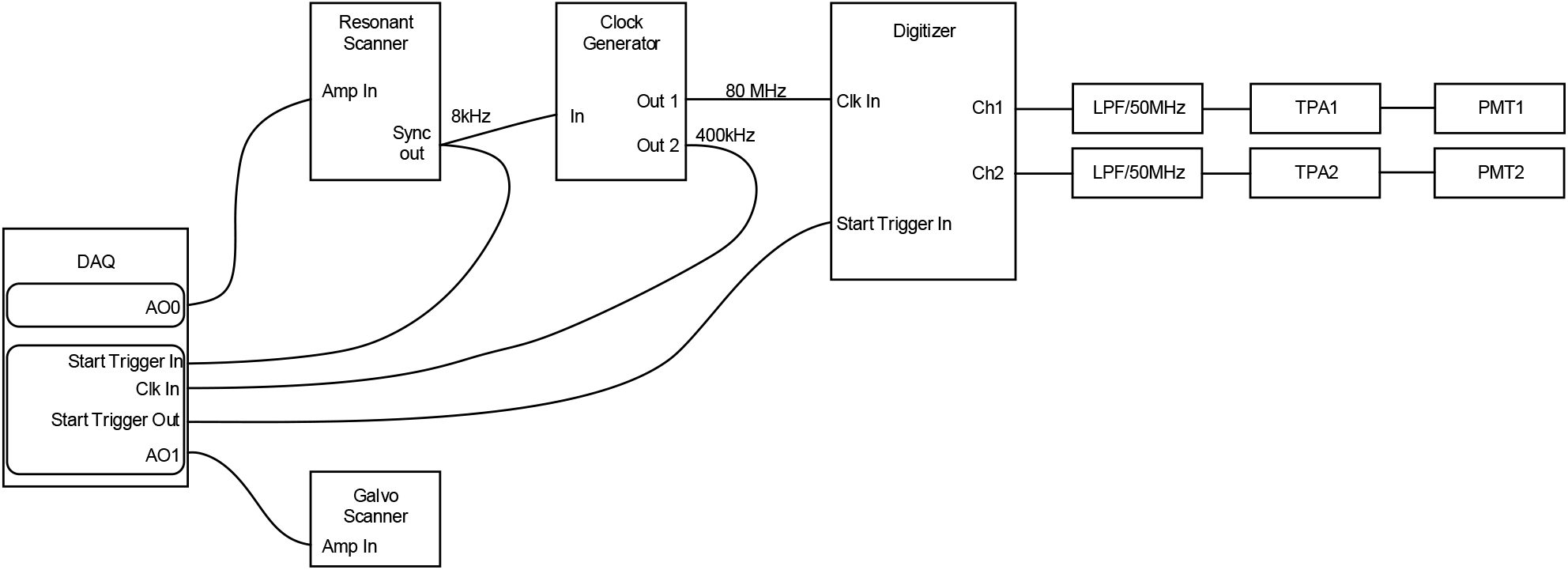
Wiring diagram for kilohertz imaging. All components are listed in Table 1.

**Supplementary Figure 3.**
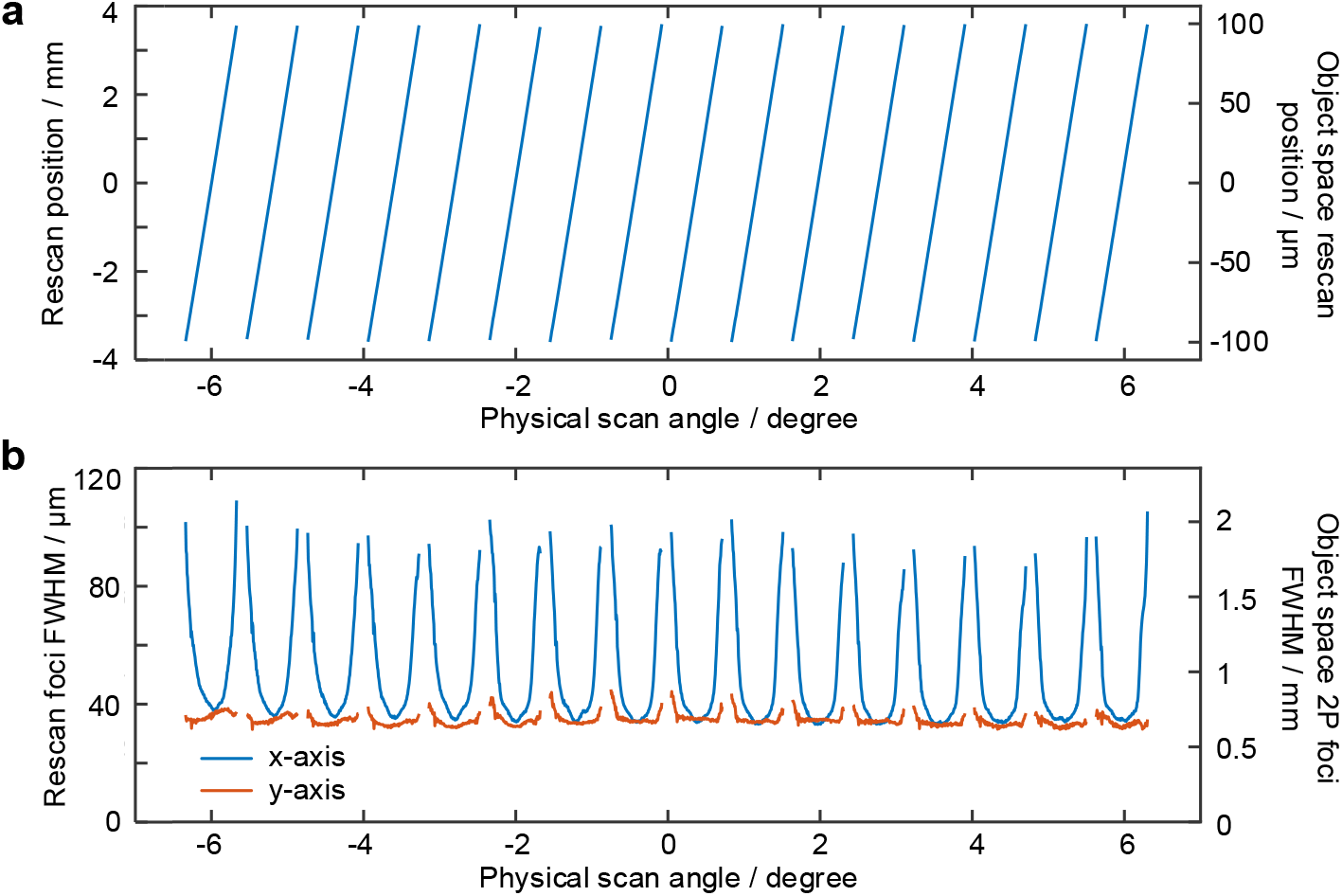
Measured rescan foci after the scan lens *f*_3_ obtained with linear galvanometer and imaged with a camera. (a) Rescan position as a function of physical scan angle. (b) FWHM of the rescan foci measured by the camera (left axis). Equivalent 2P FWHMs projected into object space were 0.97 ± 0.36 μm and 0.67 ± 0.04 μm (mean ± S.D) along the fast- and slow-axes, obtained by squaring the camera-measured intensity profiles and accounting for system magnification.

**Supplementary Figure 4.**
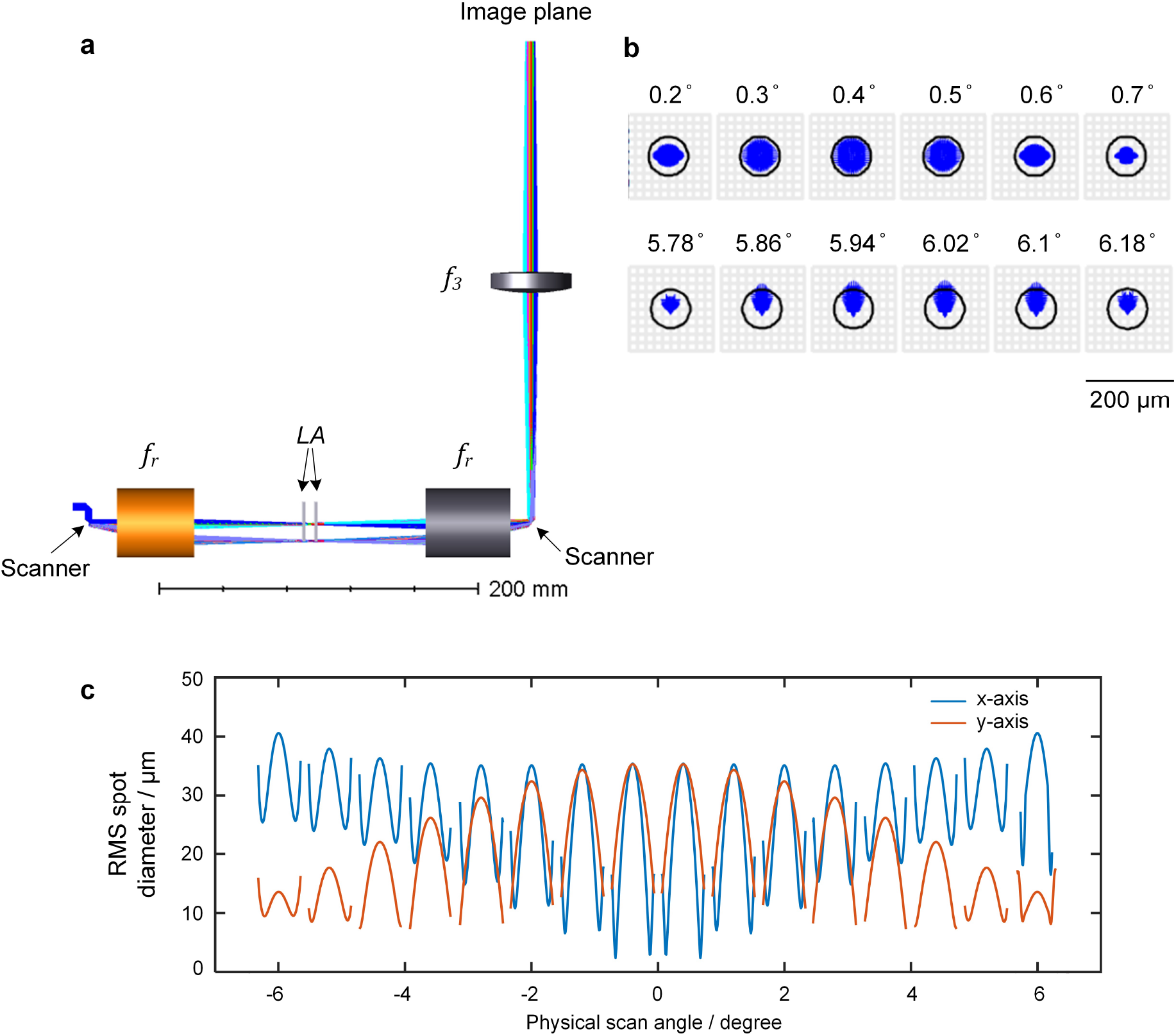
Zemax simulation of SMU1. (a) Zemax shaded model. All optical components correspond to the same labels in Fig. 1 and Table 1. The input beam was set to 3.6 mm diameter and 920 nm wavelength. For easier modeling, the SMU was unfolded about the retroflecting mirror. *f_r_* was modeled using the Zemax blackbox file of Thorlabs LSM54-850. (b) Zemax spot diagram at different scanner scan angles when the beam is transmitting through the innermost (first row, 0.2°to 0.7°) and outermost (second row, 5.78°to 6.18°) lenslet. (c) RMS spot diameter as a function of physical scan angle along the x and y axes.

**Supplementary Figure 5.**
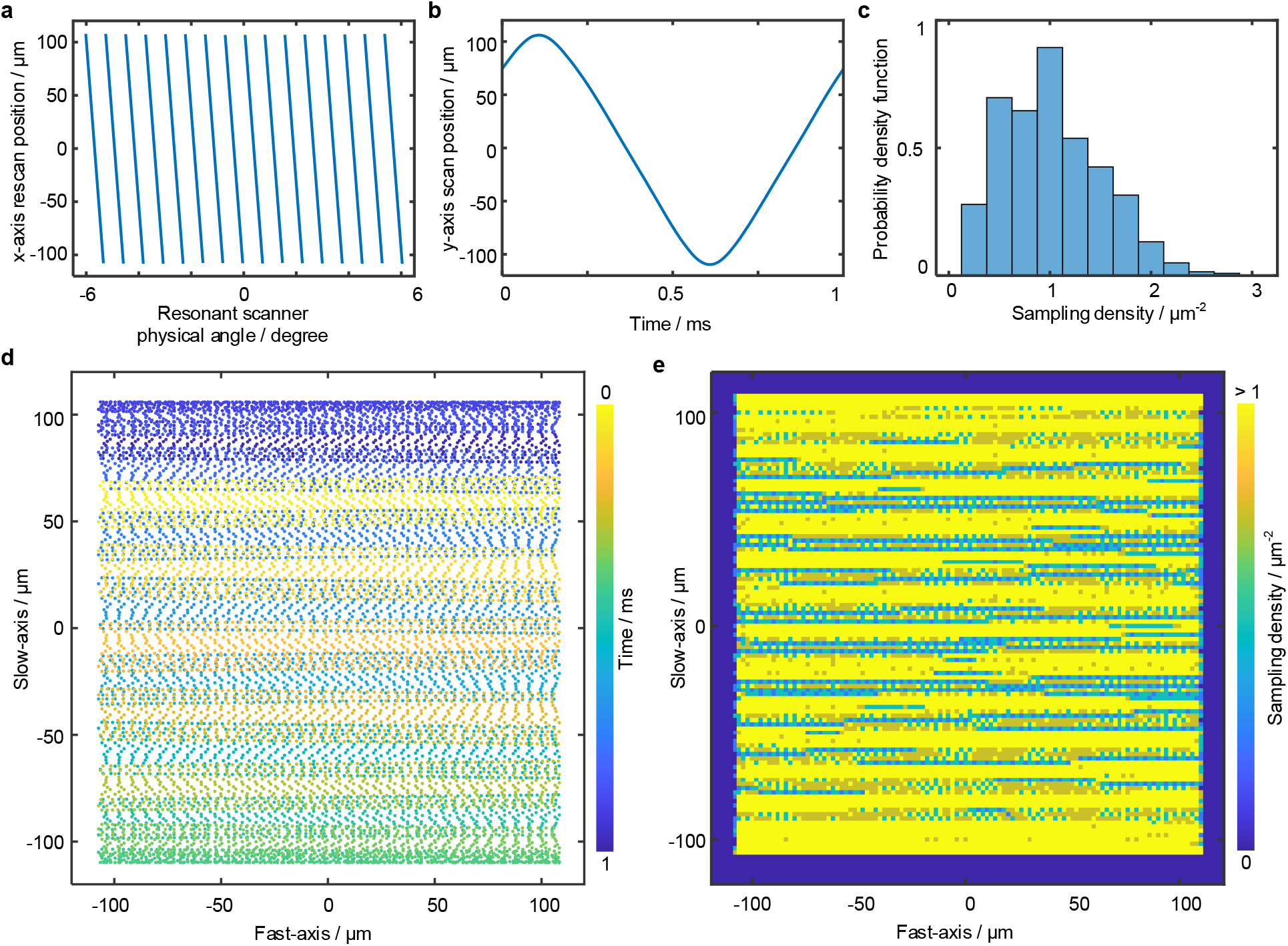
Laser pulse sampling across the imaging FOV. (a) Measured fast-axis rescan positions during a single resonant scanner scan period. (b) Measured slow-axis galvanometric scanner scan position during a single frame period. (c) Probability density function of the laser pulse sampling density (number of laser pulses per unit μm^2^) over the 200 × 200 μm FOV. The average sampling density is 1.03 ± 0.49 (mean ± std.) laser pulse/μm^2^. (d) Visualization of laser pulse sampling positions for each frame, downsampled to 20% of the original density. (e) Laser pulse sampling density across the imaging FOV.

**Supplementary Figure 6.**
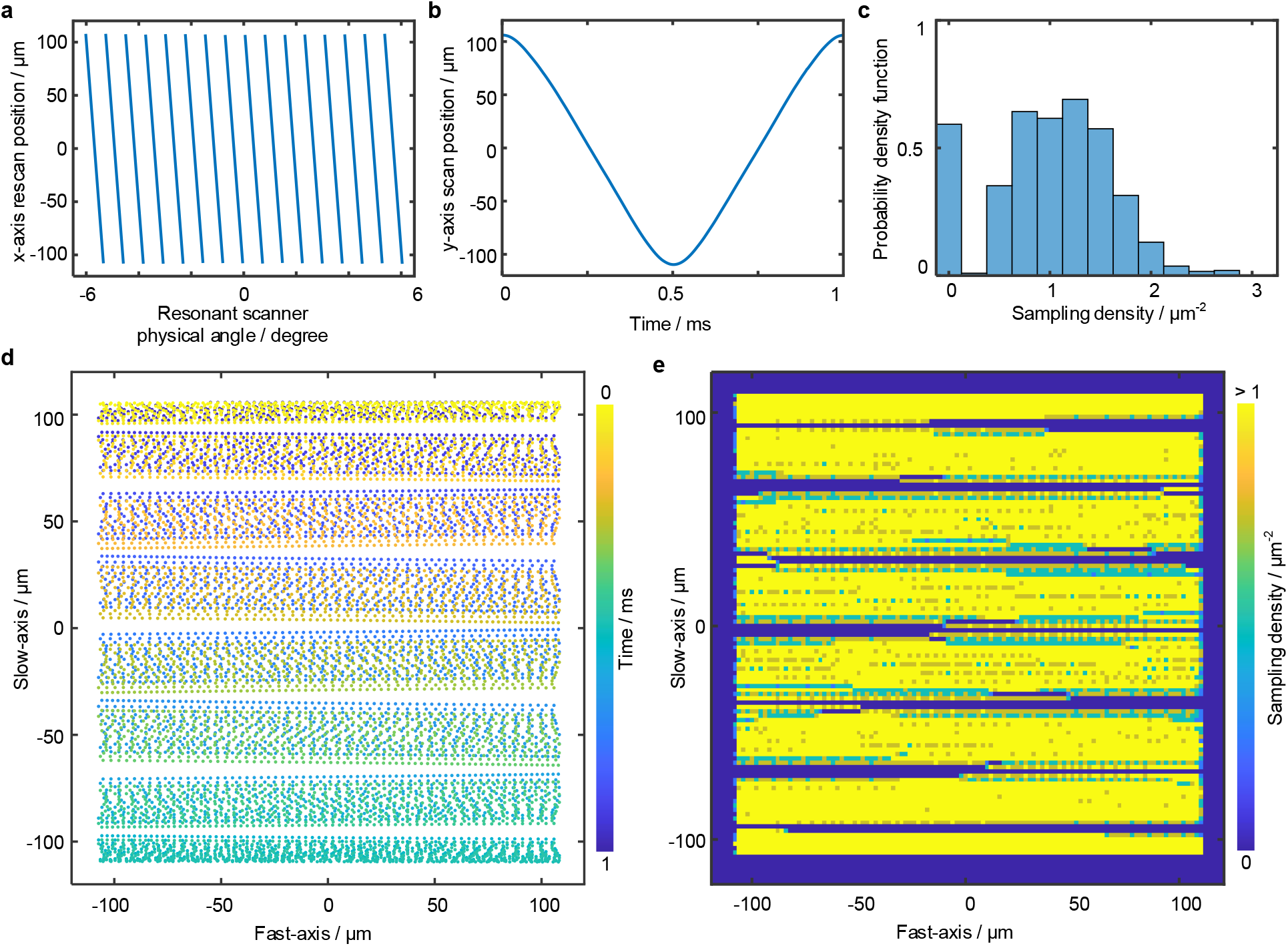
Nonuniform laser pulse sampling due to non-optimal slow-axis scanner phase. Panels in (a-e) the same as in Supplementary Fig. 5(a-e), except with the phase of the slow-axis scanner position adjusted as shown in (b). Non-optimal slow-axis scanner phase leads to significant sampling non-uniformity across the FOV, where many regions within the FOV receive no laser pulses (c,e).

**Supplementary Figure 7.**
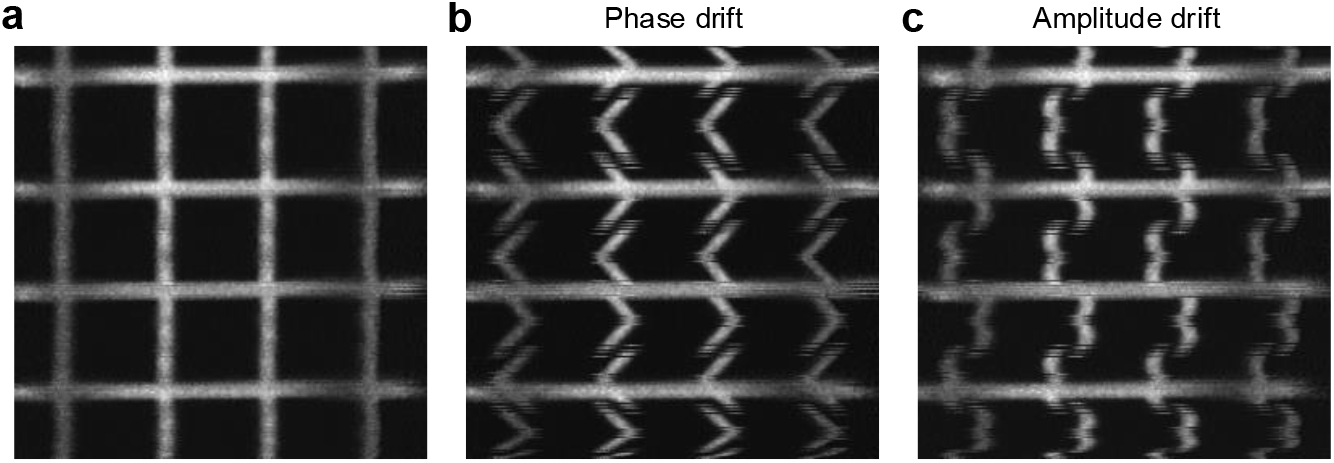
Example of image distortions due to calibration errors. (a) Reconstructed image from corrected calibration. Sample was a fluorescent grid pattern with 50 μm spacing (DA113, II-VI Aerospace Defense). (b) Reconstructed image with incorrect resonant scanner phase. (c) Reconstructed image with incorrect resonant scanner amplitude.

**Supplementary Figure 8.**
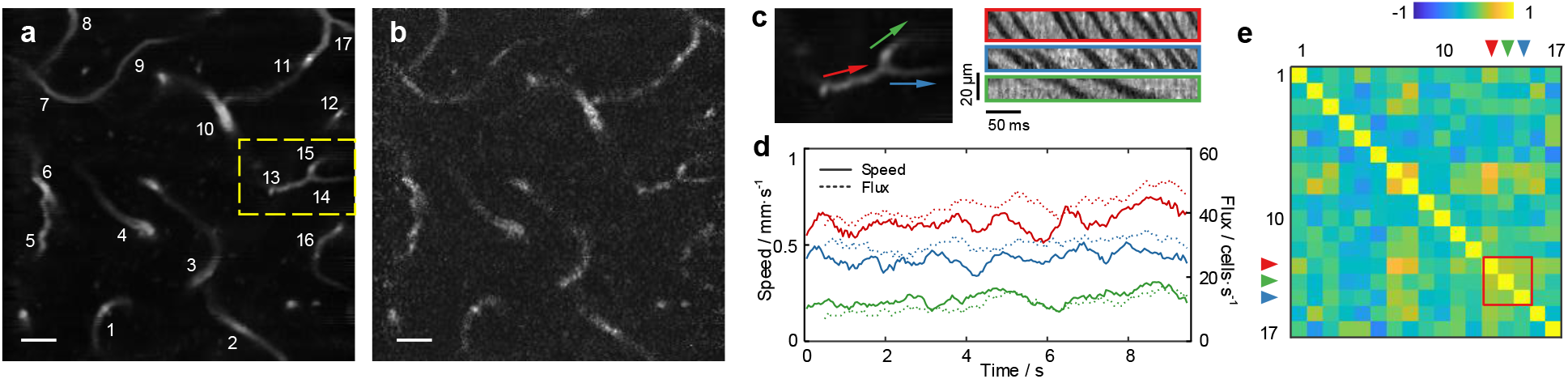
SMU-2PM enables deep tissue imaging of capillary network in vivo. (a) Averaged fluorescence image of capillary network containing 17 vessel segments at depth 340 μm. (b) Single raw frame acquired at 1 kHz. (c) Zoomed-in image (left panel) and corresponding kymographs (right panel) at a capillary junction in the yellow box region in (a). (d) Measured flow speeds and flux rates of the 3 capillary segments shown in (c). (e) Cross-correlation coefficients of the velocity profiles over the 17 capillary segments labeled in (a). Imaging depth 320 μm, excitation power 160 mW, recording duration 10 s. Scale bars in (a,b) are 20 μm.

**Supplementary Figure 9.**
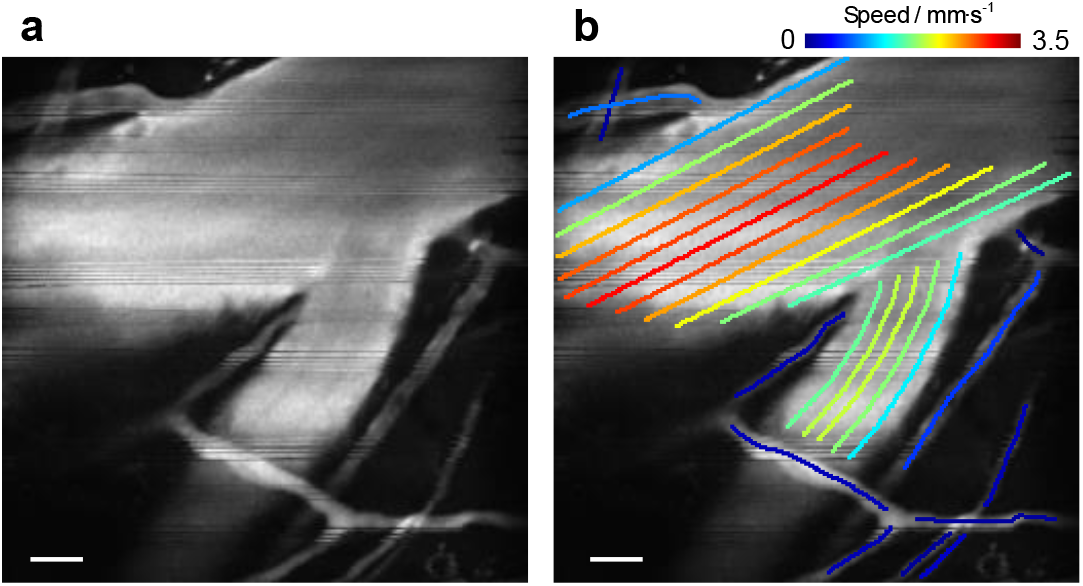
SMU-2PM enables characterization of spatially varying flow speed in large vessels. (a) Averaged fluorescence image containing a large vessel junction and multiple smaller capillaries. (b) Same as (a) but overlaid with flow speeds averaged over a 10 s recording. Imaging depth approximately 50 μm, excitation power 50 mW. Scale bars in (a,b) are 20 μm.

**Supplementary Figure 10.**
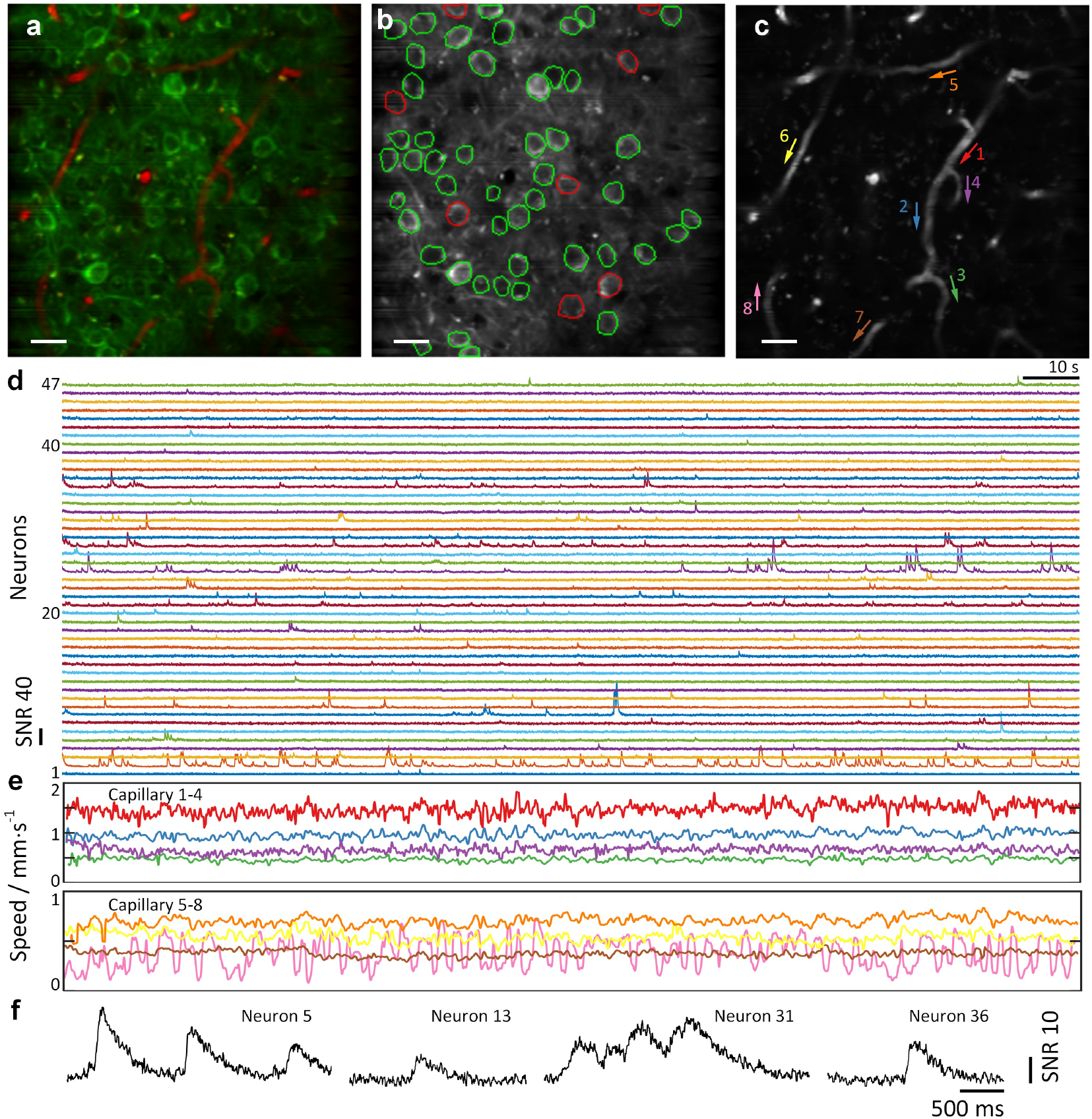
SMU-2PM enables high-throughput imaging of more than 50 neurons and 8 capillary segments in vivo. (a) Composite image of both GCaMP (green) and Texas Red (red) fluorescence. Imaging depth 150 μm, excitation power 100 mW. Recording duration 3 minutes. No stimulus was given. (b) Averaged GCaMP fluorescence image. 47 neurons that were active during the recording are outlined in green. 8 inactive neurons are outlined in red. (c) Averaged Texas Red fluorescence image with 8 capillary segments labeled with their flow directions. (d) Calcium traces of 47 neurons active during a 3 min recording. (e) Flow speed of 8 capillary segments during the same recording period, temporally aligned with (d). (f) Zoomed-in fluorescence traces from 4 selected neurons.

**Supplementary Figure 11.**
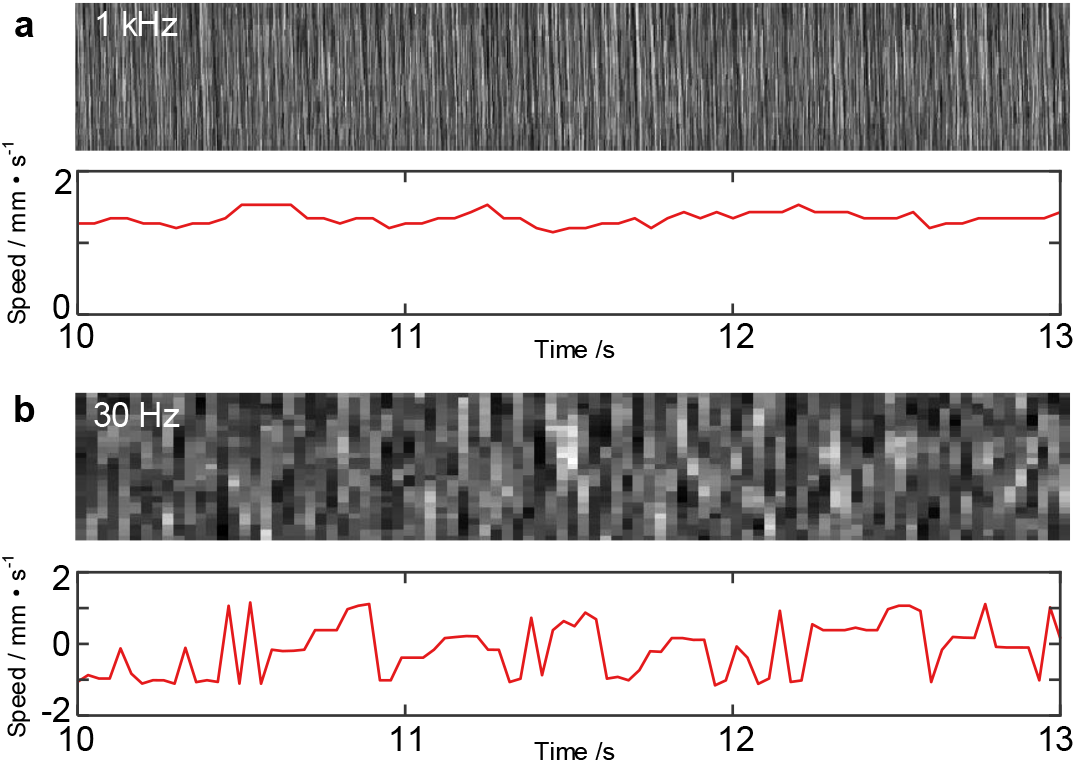
Capillary flow speed quantification at 1 kHz and 30 Hz frame rates. (a) Top panel: kymograph of capillary 1 from Supplementary Fig. 10(c) during recording period 10-13 s. Bottom panel: corresponding flow speed calculated using iterative Radon transform. (b) Same as (a) but with kymograph downsampled to 30 Hz by selecting 1 out of every 33 frames.

**Supplementary Figure 12.**
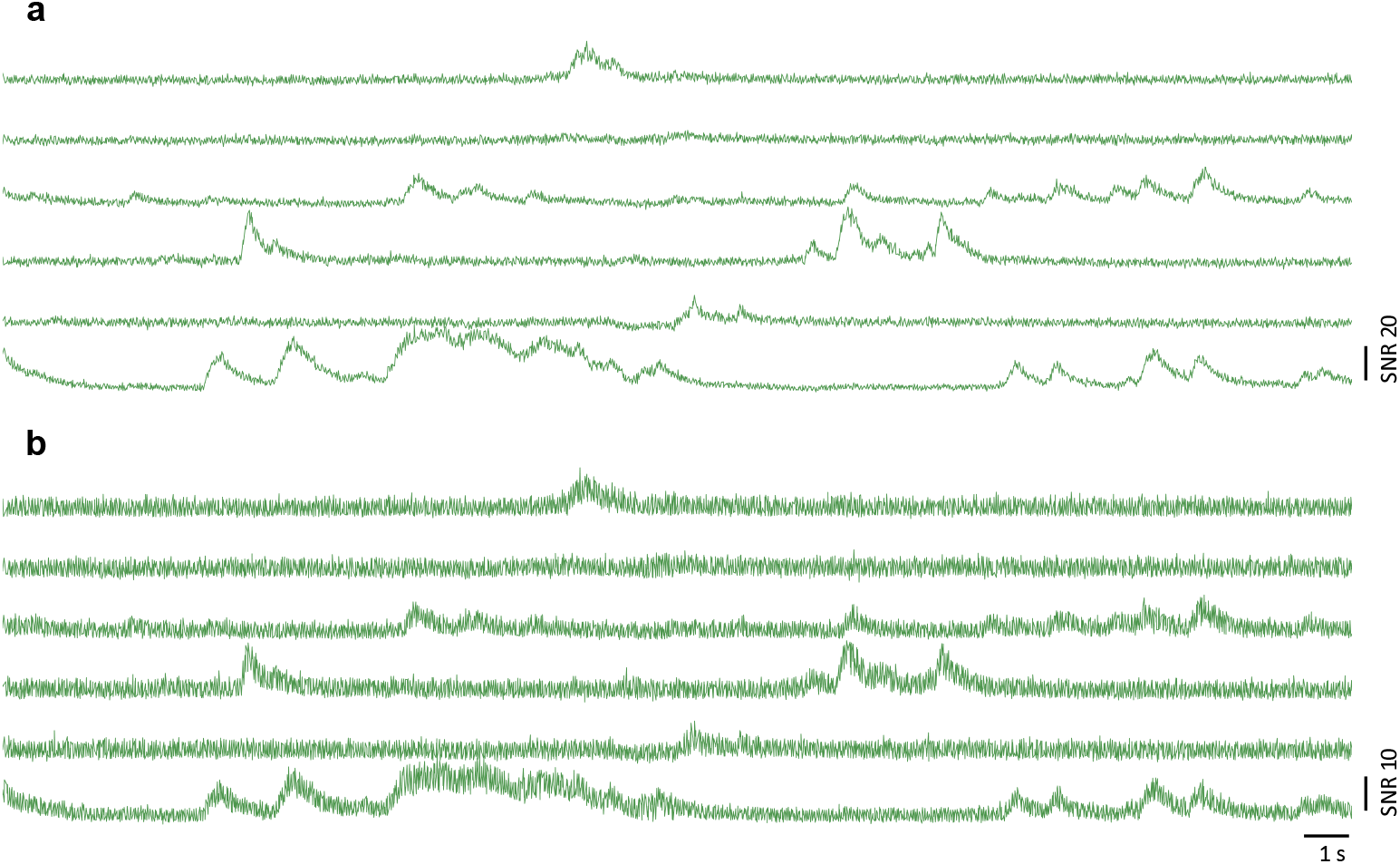
Comparison of calcium traces with and without median filter. (a) Calcium traces median filtered with 10 ms time window, corresponding to the traces shown in main Fig. 3(e) at depth 540 μm. (b) Raw calcium traces recorded at 1 kHz without median filtering.

**Supplementary Table 1.**
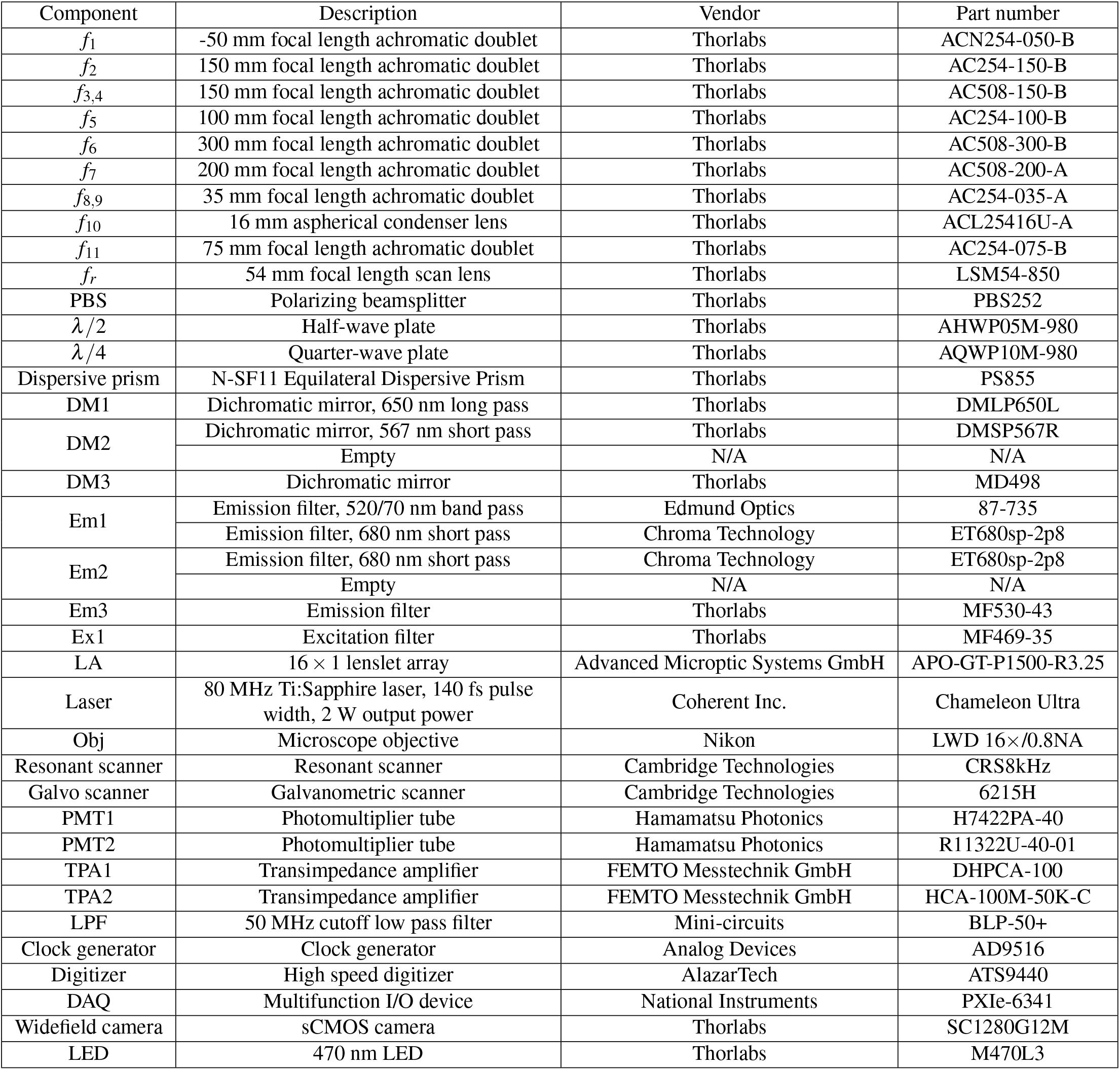
List of components.

